# Age-related differences in neurometabolite concentrations during early infancy: A cross-sectional analysis of the HBCD Release 1.0

**DOI:** 10.64898/2026.09.01.748552

**Authors:** Saipavitra Murali-Manohar, Helge J. Zöllner, Christopher W. Davies-Jenkins, Aaron T. Gudmundson, Steve C. N. Hui, Yulu Song, Gizeaddis L. Simegn, Zahra Shams, Abdelrahman Gad, Borjan Gagoski, M. Dylan Tisdall, Muhammad G. Saleh, Ralph Noeske, Sandeep K. Ganji, Guillaume Gilbert, Yansong Zhao, Timothy J. Hendrickson, Erik G. Lee, William T. Clarke, Douglas C. Dean, Mary Beth Nebel, Christopher D. Smyser, Damien A. Fair, Heather E. Volk, Georg Oeltzschner, Jessica L. Wisnowski, Richard A. E. Edden, the HBCD MRS Working Group

## Abstract

**Background:** Developmental trajectories of low-concentration neurometabolites such as the neurotransmitter γ-aminobutyric acid (GABA), and the antioxidants glutathione (GSH) and ascorbate (Asc) across early infancy remain unexplored. Advances in spectral editing enabled the measurement of these key molecules together with high-concentration metabolites like N-acetylaspartate (NAA) and glutamate (Glu) in the HEALthy Brain and Child Development (HBCD) study, the largest longitudinal study of early brain development in the United States.

**Purpose:** To determine the age-associated trajectories of 14 key neurometabolites during early infancy from a cross-sectional ^1^H-MRS dataset.

**Materials and Methods:** HBCD utilizes ISTHMUS, an integrated MRS sequence that includes both an unedited short-echo-time PRESS acquisition and an advanced 4-step Hadamard-encoded sequence, HERCULES to enable measurement of both high- and low-concentration metabolites. Metabolite quantification was carried out by the HBCD Data Coordinating Center (HDCC) using an automated Osprey pipeline, with data from 201 infants ages 0 – 10 weeks adjusted age included in the tabular imaging results in HBCD data release 1.0. After excluding preterm born infants and data with poor linewidth or model quality metric, we tested for linear associations with adjusted age for each metabolite.

**Results:** Concentration estimates of total NAA (tNAA), total creatine (tCr) and glutamate (Glu) as well as the combined sum of glutamate and glutamine (Glx) significantly increased across ages 0 – 10 weeks, while myo-inositol (mI) decreased. GABA and GSH showed age-related trends, but did not reach significance. Levels of the lipid precursor phosphorylethanolamine (PE) and Asc are higher in the first months than established adult values.

**Conclusion:** Multiple metabolites showed significant age-related changes during early infancy. While GABA and GSH did not, future work will establish whether the trends suggested here contribute to linear or non-linear patterns across the first years of life.

**Summary Statement:** HBCD MRS Data Release 1.0 cross-sectional analysis of tabulated results shows significant changes of neurometabolite levels in the thalamus from birth to 10-week-old infants.

**Key Results:** This study provides the first ever concentration level reports of multiple low-concentration metabolites such as GABA+, GSH, PE, Lac, Asc as well as their cross-sectional trajectory in early postnatal period of brain development.

## Introduction

Infant brain development is characterized by rapid brain growth and myelination (1), which is accompanied by maturational changes in energy metabolism, neurotransmission and antioxidant defense systems. Magnetic resonance spectroscopy (MRS), a non-invasive technique, offers unprecedented insight into cell-specific neurochemical processes (2). Although increasingly applied across the adult lifespan, few in vivo MRS studies have explored the dynamic neurochemical changes that underlie early development. Measuring neurochemistry in early infancy will provide critical insights not only into healthy brain development, but also how alterations in this early neurochemistry contributes to adverse neurodevelopmental outcomes.

The knowledge gap in pediatric MRS stems from several challenges: lack of robust acquisition techniques; long acquisition durations to measure lower-concentration metabolites; a paucity of age-appropriate relaxation reference values; reduced subject compliance; and inaccessibility of motion-correction methods. The existing limited pediatric MRS literature clearly indicates that neurochemical concentration levels change during healthy brain development (3– 5). The trajectories of easily-detectable, high-concentration neurometabolites, such as total N-acetyl aspartate (tNAA), glutamate (Glu), total creatine (tCr), total choline (tCho), myo-inositol (mI), taurine (Tau), show rapid change in the first three months of life (4).

However, low-concentration J-coupled neurometabolites, including γ-aminobutyric acid (GABA), glutathione (GSH), lactate (Lac), phosphorylethanolamine (PE), and ascorbate (Asc), have never been explored in early infancy. These metabolite signals are challenging to reliably measure and quantify at clinical field strengths due to overlapped J-coupled peaks. Robust measurement of these metabolite signals at 3T, requires advanced spectral editing techniques (6). HERCULES editing scheme (7), enables simultaneous measurement of multiple J-coupled metabolite signals in a single acquisition. This drastically reduces total scan durations, improving feasibility for pediatric MRS.

The HEALthy Brain and Child Development (HBCD) study is the largest longitudinal study of early brain and child development in the United States with 26 recruiting sites across the nation (8). The primary objective of the study is to understand normal brain development and elucidate how early environments shape the trajectory from gestation through the first decade of life. HBCD utilizes a novel multi-sequence, ISTHMUS (8, 9) (Figure 1A), which combines short-TE PRESS, HERCULES (10) editing, and dual-TE interleaved water reference scans to mitigate frequency drift in a long MRI protocol. Moreover, the PRESS data are collected at the beginning, maximizing data collection during natural sleep, when infants may awaken prematurely before the end of the sequence. HBCD is designed to be a scientific resource, with annual data releases made available through a controlled-access repository. This study investigates the cross-sectional linear effects of age on metabolite levels from the bilateral thalamus (Figure 1B) during the first ten weeks of life, using tabulated data from HBCD Data Release 1.0.

**Figure 1:**
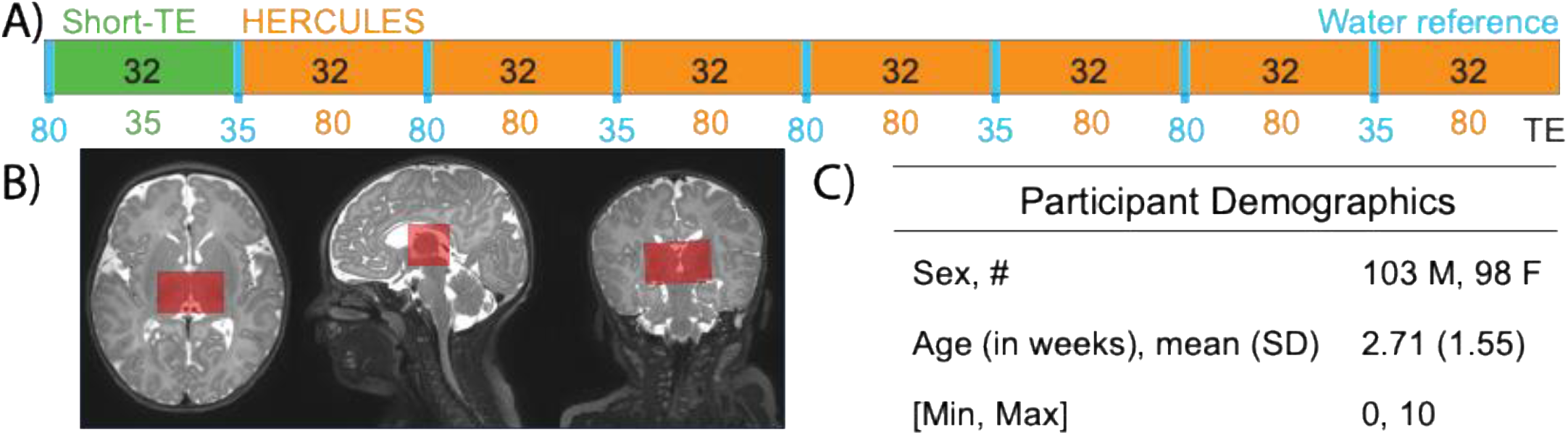
**A)** ISTHMUS sequence combining short-TE unedited PRESS and HERCULES used in the HBCD study MRS data acquisition **B)** Voxel placement in the bilateral thalamus of an infant **C)** Visit-2 (0–1 month) participant demographics for MRS data from the tabulated results of HBCD Data Release 1.0. Here, ‘Age’ refers to adjusted age, as released by HBCD, which is estimated date of delivery to the measurement date rounded down to whole weeks.

## Methods

### HBCD Data Release 1.0

The ongoing HBCD study aims to recruit over 6000 mother-child pairs and to follow their children across the first decade of life. The full protocol has been described elsewhere (8). Recruitment began in July 2023 and is ongoing. The first data release (HBCD 1.0, available June 2025 on the NIH Brain Development Cohorts (NBDC) Data Repository), includes all data collected through June 2024.

Release 1.0 includes pre-processed MRS data from 201 newborn infants at Visit 2 and 14 infants ages 3 – 8 months at Visit-3. Per regulations, data were downloaded into an NIST-SP-800-171-compliant computing environment following approval of the HBCD Data Use Certification and successful completion of responsible use of data training. For this analysis, only Visit-2 participants were considered (Figure 1C) given the limited number of Visit-3 datapoints. HBCD reports ‘adjusted age’ calculated as the time from the expected due date to the date of measurement rounded down to whole weeks, spanning 0–10 weeks for Visit-2. This is analogous to 40 – 50 weeks, measured as post-conceptional age (PCA).

### MRS Data Acquisition

HBCD is carried out using a single protocol harmonized across 26 participating sites, with 19 sites operating Siemens 3T Prisma, 2 operating GE 3T MR750s, 2 operating Philips 3T MR7700s, 2 operating Philips 3T Achieva dStream MR scanners and 1 operating a 3T Philips Elition RX scanner(8). Written informed consent was obtained from the parent or legal guardian in accordance with centralized IRB approval. The Visit-2 MRI protocol includes structural T_2_-weighted MRI (resolution: 0.8×0.8×0.8 mm), functional (fMRI), diffusion (dMRI), QALAS (11), ISTHMUS (9), and structural T_1_-weighted MRI (8). The protocol is quasi-randomized across participants, with T_2_-weighted MRI acquired first, followed by fMRI and dMRI, in random order, then QALAS(11) and ISTHMUS, in random order, and finally T_1_-weighted MRI. Since data are collected during natural sleep, some infants awoke early and do not complete the full protocol.

Lower resolution axial and coronal T_2_-weighted MRI scans preceded the MRS acquisition, and were used as MRS localizers facilitating voxel placement and co-registration to T_2_-weighted MRI(8). A 23×30(RL)×23 mm^3^ voxel was placed in the bilateral thalamus (Figure 1B). ISTHMUS (Figure 1A, Supplementary Table 1 (12)) data were acquired with 32 short-TE PRESS (TE/TR: 35/2000 ms), 224 HERCULES transients (TE/TR: 80/2000 ms) and interleaved short- and long-TE water reference scans (4 transients each).

**Table 1:**
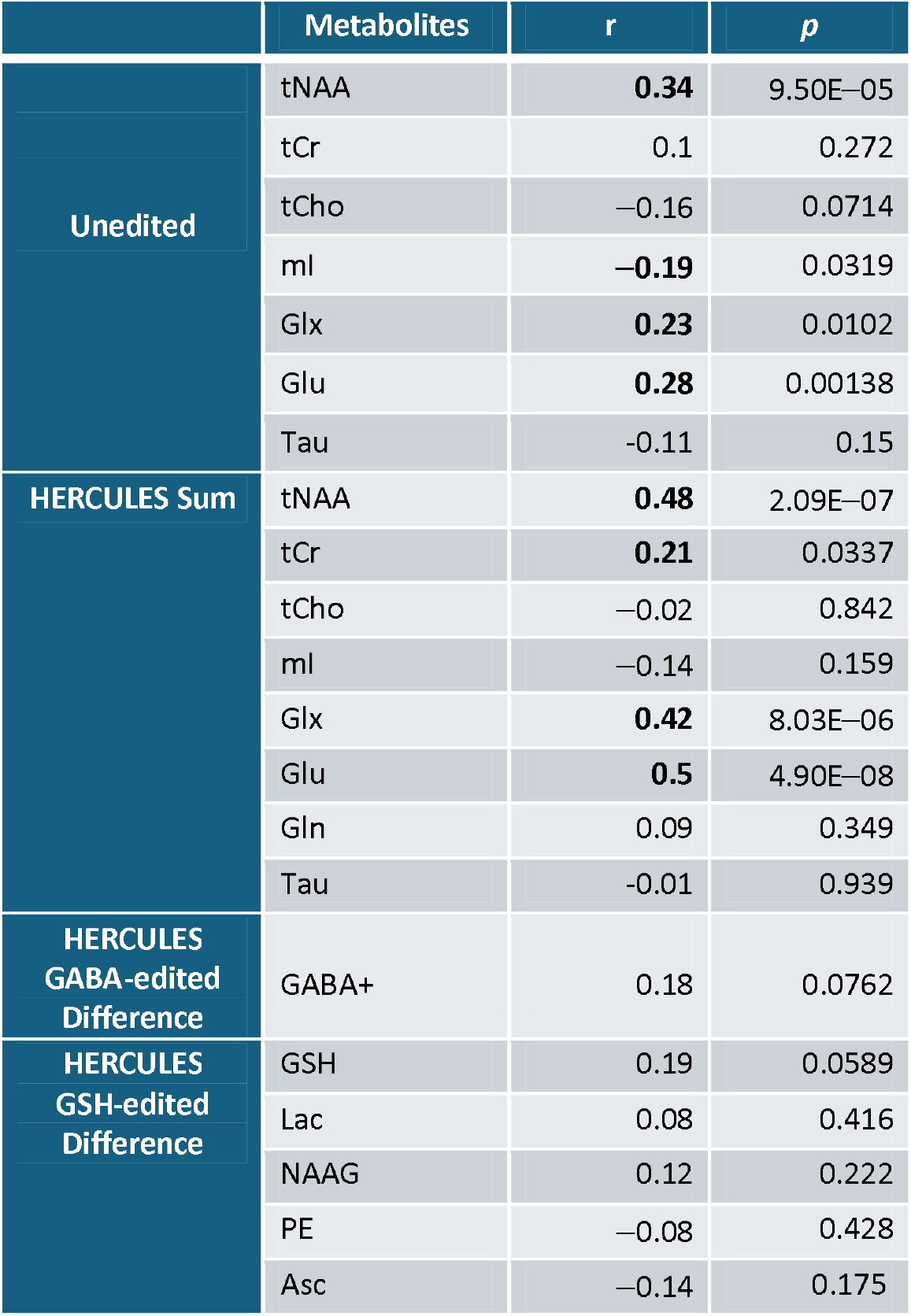
Pearson correlation coefficients between metabolite concentration estimates and adjusted age 0–10 weeks, along with the respective *p*-values. Bolded r indicated significant correlations (*p* < 0.05).

| | Metabolites | $r$ | $p$ |
| --- | --- | --- | --- |
| Unedited | tNAA | **0.34** | 9.50E–05 |
|  | tCr | 0.1 | 0.272 |
|  | tCho | –0.16 | 0.0714 |
|  | ml | **–0.19** | 0.0319 |
|  | Glx | **0.23** | 0.0102 |
|  | Glu | **0.28** | 0.00138 |
|  | Tau | –0.11 | 0.15 |
| HERCULES Sum | tNAA | **0.48** | 2.09E–07 |
|  | tCr | **0.21** | 0.0337 |
|  | tCho | –0.02 | 0.842 |
|  | ml | –0.14 | 0.159 |
|  | Glx | **0.42** | 8.03E–06 |
|  | Glu | **0.5** | 4.90E–08 |
|  | Gln | 0.09 | 0.349 |
|  | Tau | –0.01 | 0.939 |
| HERCULES GABA-edited Difference | GABA+ | 0.18 | 0.0762 |
| HERCULES GSH-edited Difference | GSH | 0.19 | 0.0589 |
|  | Lac | 0.08 | 0.416 |
|  | NAAG | 0.12 | 0.222 |
|  | PE | –0.08 | 0.428 |
|  | Asc | –0.14 | 0.175 |

|  | Metabolites | *r* | *p* |
| --- | --- | --- | --- |
| Unedited | tNAA | **0.34** | 9.50E–05 |
|  | tCr | 0.1 | 0.272 |
|  | tCho | –0.16 | 0.0714 |
|  | ml | **–0.19** | 0.0319 |
|  | Glx | **0.23** | 0.0102 |
|  | Glu | **0.28** | 0.00138 |
|  | Tau | –0.11 | 0.15 |
| HERCULES Sum | tNAA | **0.48** | 2.09E–07 |
|  | tCr | **0.21** | 0.0337 |
|  | tCho | –0.02 | 0.842 |
|  | ml | –0.14 | 0.159 |
|  | Glx | **0.42** | 8.03E–06 |
|  | Glu | **0.5** | 4.90E–08 |
|  | Gln | 0.09 | 0.349 |
|  | Tau | –0.01 | 0.939 |
| HERCULES GABA-edited Difference | GABA+ | 0.18 | 0.0762 |
| HERCULES GSH-edited Difference | GSH | 0.19 | 0.0589 |
|  | Lac | 0.08 | 0.416 |
|  | NAAG | 0.12 | 0.222 |
|  | PE | –0.08 | 0.428 |
|  | Asc | –0.14 | 0.175 |

### Data Processing and Quantification

Metabolite quantitation was carried out at the HBCD Data Coordinating Center (8), using an automated (13) Osprey 2.8.1 (7) pipeline and overseen by the HBCD MRS Working Group. Multi-vendor raw MRS data were converted to NIfTI-MRS(14) and organized in BIDS (15). Following this, all MRS analyses were performed in Osprey(7) for PRESS and HERCULES spectra (Supplementary Table 2 (12)). Pre-processing of MRS data involves receiver coil combination, alignment of individual transients using robust spectral registration(16), eddy-current correction, and coherent averaging to produce short-TE spectra as well as GABA-edited, GSH-edited, and Sum spectra from HERCULES. Binary MRS voxel masks were generated in the MRS localizer space and then co-registered with the structural images. Brain segmentation of T_2_ images was performed using BIBSNet (17,18). BIDS App 2.4.3 (https://hub.docker.com/r/dcanumn/osprey-bids/tags) was used to support co-registration between the structural images and the MRS localizers. Infants that had both PRESS and HERCULES data and successful segmentation were included in the tabulated results in the HBCD Data Release 1.0, with future releases expected to include participants with only partial data. Water-scaled tissue and relaxation-corrected metabolite concentration estimation (19) was carried out using the Osprey default values for tissue-specific water densities, water and metabolite relaxation times.

The analysis procedure separately models the short-TE, HERCULES-Sum, GABA-edited difference and GSH-edited difference spectra with density-matrix-simulated metabolite basis functions, Gaussian basis functions for macromolecules (MM)/lipids, and a cubic-spline baseline. Co-edited MM in the GABA-edited spectrum was modeled with a Gaussian function, and the composite GABA+MM_3.0_ (GABA+) was used in analyses. The tabulated results include data quality metrics: tCr signal-to-noise ratio (SNR); tCr linewidth (at full-width half-maximum); water linewidth; residual water amplitude; pre-processing frequency shift; and model relative residual (relRes), i.e., normalized sum of squares of the residual divided by the variance of the spectral noise(20). This analysis considers: tNAA, tCr, tCho, mI, Glx, Glu, and Tau levels from short-TE models; tNAA, tCr, tCho, mI, Glu, Gln, Glx, and Tau levels from HERCULES-Sum; GABA+ levels from GABA-difference models; GSH, PE, NAAG, Asc, and Lac from GSH-difference models. Water-referenced concentration values were used through the img_osprey_*sequence* _A_TissCorrWaterScaled_Voxel_1_Basis_1_*met* variables in the output tables, where *met* stands for the respective metabolite and *sequence* is either unedited, HERCULES_sum, HERCULES_diff1, or HERCULES_diff2 for the short-TE, HERCULES-Sum, and GABA- and GSH-edited difference spectra respectively. All four models used the shorter-TE water reference for quantification, with reduced T_2_-weighting, and TE-matched data for eddy-current correction.

### Quality Assurance and Exclusion of Data

To maximize the reliability and reproducibility of our results, we considered both subject factors and data quality in generating the final dataset. Of the 201 infants with Visit-2 MRS, 13 were born preterm (gestational age at delivery < 37 weeks). These were excluded from these analyses due to previous studies documenting an effect of preterm birth on neurometabolism (21). On data quality, the HDCC processing pipeline flags data with tCr linewidth > 12 Hz; however, this does not reliability identify every dataset with poor fit. Thus, while we continue to refine automated pipelines for identifying low quality data in large-scale datasets like HBCD, we chose to apply a highly conservative threshold for data quality here in the first analysis of HBCD MRS data. Specifically, data were excluded from these analysis if: : tCr linewidth > 5 Hz; and with outliers (more than one inter-quartile range from the inter-quartile limits) based on spectral relative residual (relRes) for PRESS and HERCULES-Sum spectra. This conservative approach, compared to community norms (12) (tCr linewidth < 12 Hz), was applied here while we continue to validate automated quality control processes specific to pediatric MRS data. Pearson correlation analysis was conducted in MATLAB R2024a to test the relationship between adjusted age and tissue-corrected water-scaled metabolite levels in institutional units. Linear correlations were assessed using Pearson’s correlation coefficient r. –1 < r < 1, where –1 shows perfect negative correlation, +1 perfect positive correlation and 0 indicates no correlation. No correction of *p*-values is applied to account for multiple comparisons. Correlations with *p* < 0.05 were considered significant.

## Results

Before applying exclusion criteria for MRS data, mean (± standard deviation) tCr linewidth and SNR for the tabulated Visit-2 results were 4.8 ± 2.4 Hz, 80 ± 24 for PRESS spectra and 5.1 ± 2.5 Hz, 145 ± 53 for HERCULES spectra, respectively. Histograms of these data quality metrics (before exclusion) are shown in Figure 2A. We present the STROBE diagram in Figure 2B showing application of all exclusion criteria and the resultant sample. After exclusion of preterm datapoints, a total of 188 Visit-2 datasets were checked for tCr linewidth > 5 Hz and relRes outliers. 42 and 52 datasets had tCr linewidth > 5 Hz for PRESS and HERCULES, respectively. relRes outliers were: 16 PRESS; 24 HERCULES Sum; 46 GABA-edited difference; and 43 GSH-edited difference spectra. Finally, the combined numbers of exclusions were: 47 PRESS; 78 HERCULES-Sum; 88 GABA-edited difference; and 85 GSH-edited difference spectra. Therefore, the total number of datasets included for statistical analysis were 141, 110, 100 and 103 for PRESS and HERCULES-Sum/GABA-edited difference/GSH-edited difference spectra.

**Figure 2:**
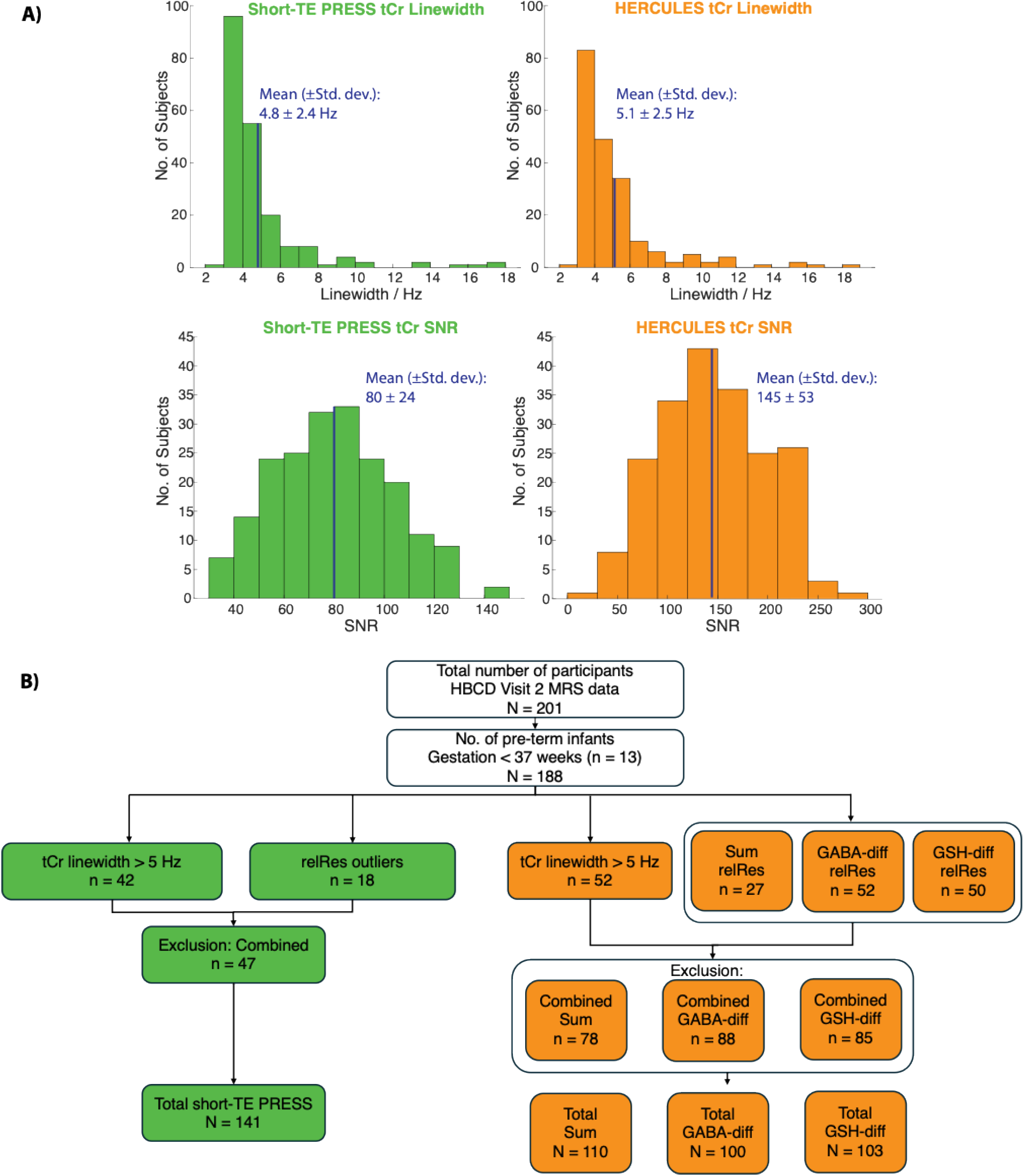
**A)** tCr SNR and tCr linwidth histograms for PRESS and HERCULES **B)** STROBE diagram of subject exclusions.

After QC, the mean tCr linewidth and SNR were 3.9 ± 0.5 Hz, 87 ± 22 for PRESS data and 3.9 ± 0.5 Hz, 165 ± 49 for HERCULES spectra respectively. Median relRes spectra, Osprey model and fit residual (representative of the Visit-2 MRS data model) are shown in Figure 3. BIBSNet segmentations were manually corrected for HBCD Data Release 1.0 and have higher quality compared to an automated run of BIBSNet (18).

**Figure 3:**
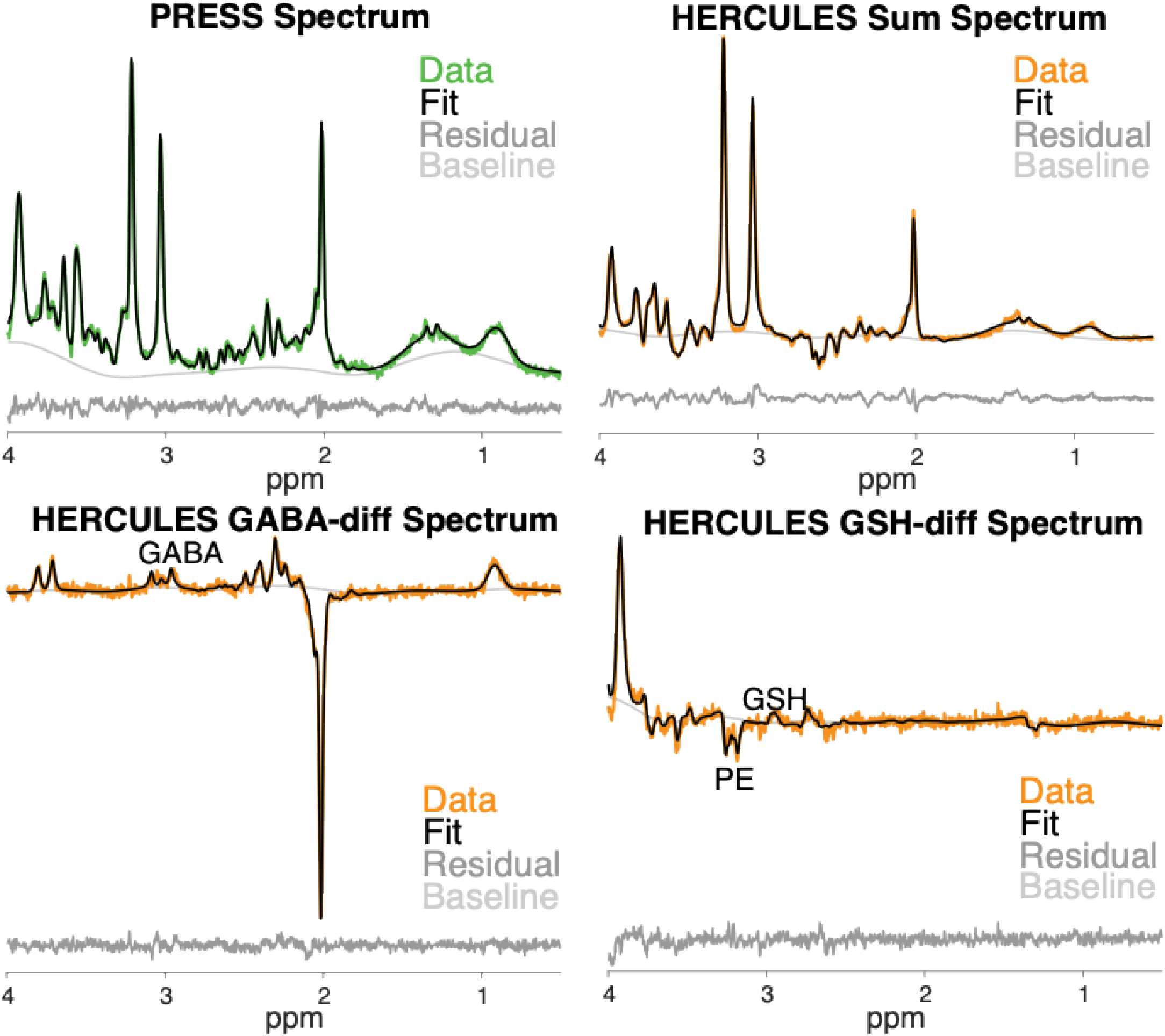
Short-TE and HERCULES-edited PRESS spectra from 2-week and 4-week adjusted-age old infants, respectively, chosen from calculated relRes medians, from short-TE PRESS (median relRes: 3.4) and HERCULES-Sum (median relRes: 9.0) modeling respectively.

We observed significant linear correlations with age across multiple metabolites from both short-TE PRESS and HERCULES data. Starting first with the results from the PRESS spectra, we observed linear increases with age for tNAA (r = 0.34), Glu (r = 0.28), and Glx (r = 0.23), the sum of Glu + Gln, and linear decreases with age for mI (r = – 0.19) (Figure 4, Table 1). There were no significant linear effects of age for tCr, tCho and Tau.

**Figure 4:**
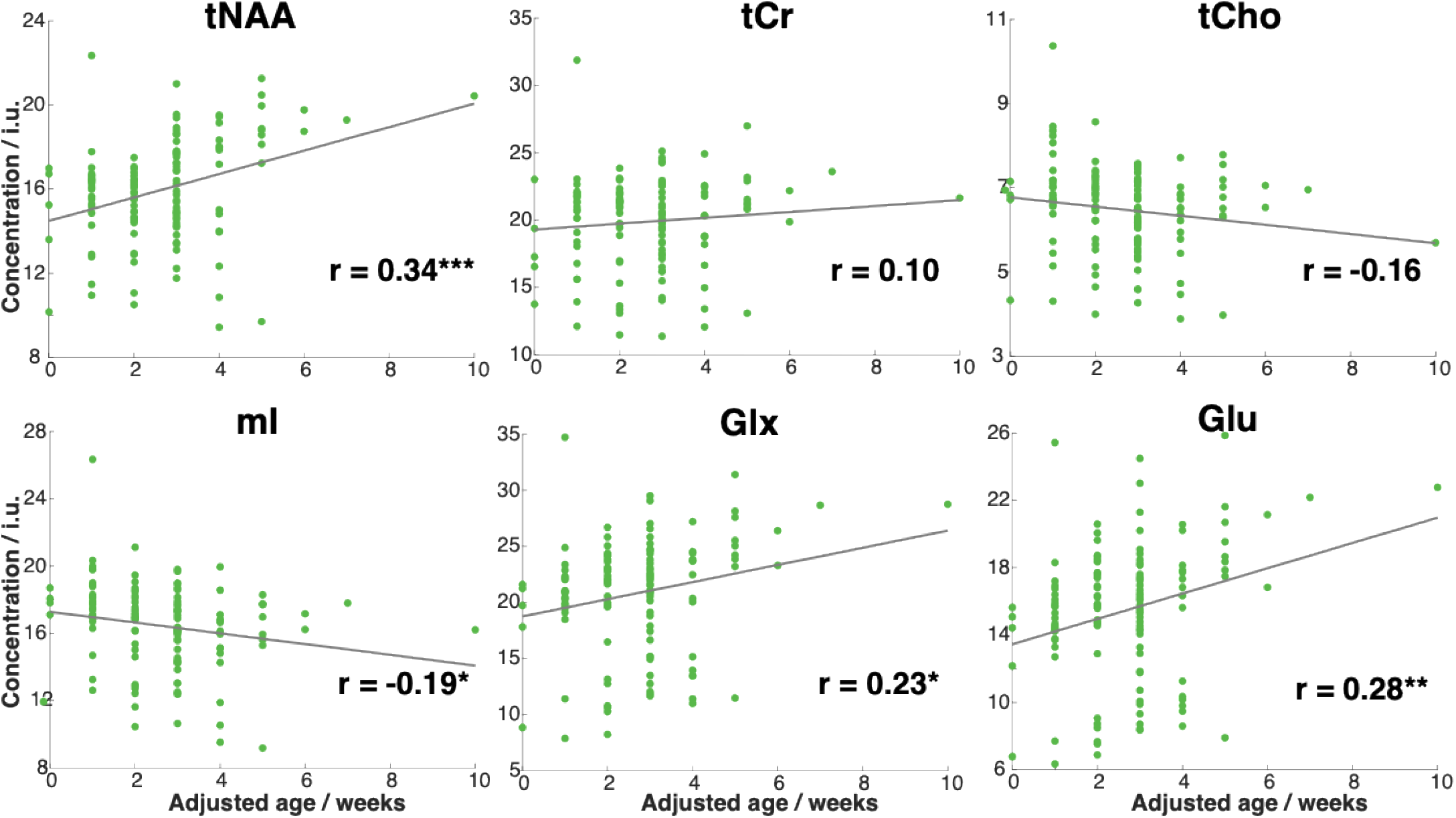
Correlation plots of adjusted age (‘Adjusted age’ 0 week corresponds to PCA 40 weeks) vs. short-TE-PRESS-derived metabolite concentration level estimations. r indicates Pearson correlation coefficient. Asterisk (*) indicates significant correlations. * *p* <0.05, ** *p*<0.01, \*\*\**p*<0.001

For HERCULES, we again observed strong linear increases with age for tNAA (r = 0.48), Glu (r = 0.50) and Glx (r = 0.42) as well as an age-related increase for tCr (r = 0.21) (Figure 5, Table 1). There was a trend toward a linear decrease with age for mI (r = –0.14, *p* = 0.16) that did not reach significance. As above, there were not significant linear effects of age on measured tCho and Tau.

**Figure 5:**
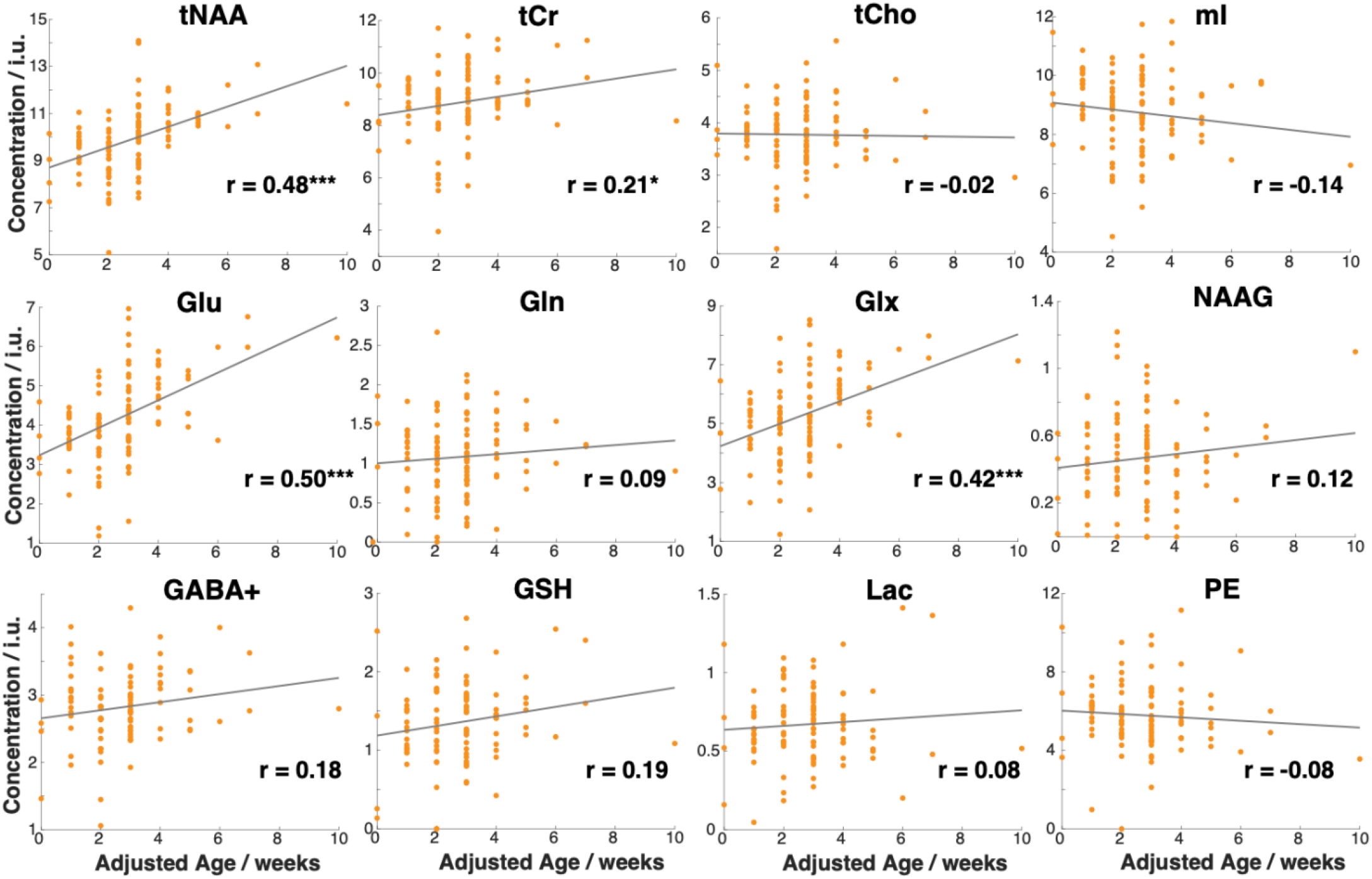
Correlation plots of adjusted age (‘Adjusted age’ 0 week corresponds to PCA 40 weeks) vs. HERCULES-derived metabolite concentration level estimations. r indicates Pearson correlation coefficient. Asterisk (*) indicates significant correlations. * *p* <0.05, ** *p*<0.01, \*\*\**p*<0.001.

Among the key low-concentration metabolites, GABA+ (r = 0.18, *p* = 0.08) and GSH (r = 0.19, *p* = 0.06) showed trends toward linear increases with age, but neither reached statistical significance in this analysis. None of the remaining low-concentration metabolites showed significant age-related effects.

Mean concentration levels (± standard deviation) in institutional units for the low-concentration metabolites measured from HERCULES are: GABA+ 2.83 ± 0.56; GSH 1.36 ± 0.52; Lac 0.67 ± 0.24; NAAG 0.47 ± 0.27; PE 5.79 ± 1.74; Asc 2.78 ± 0.89; Gln 1.08 ± 0.50.

## Discussion

The inaugural HBCD data release is a landmark dataset in MRS with 201 datasets from 0–10 week-old infants. The normative estimates from this study may provide a reference for healthy development, especially for future studies investigating deviations from typical development. The largest previous study(4) in pediatrics, focused largely on the cortical gray (GM) and white matter (WM), included relatively few data points in the deep gray nuclei (DGN) and fewer than a dozen in the neonatal period. Although neurodevelopment is inherently nonlinear (4), linear models were used as an initial approximation over the restricted early infancy age range. In our study, tNAA, Glu and Glx showed robust linear increases with age consistent with prior findings (4). NAA is synthesized in neuronal mitochondria and the acetyl groups from NAA are an important carbon source during myelination. Myelination in the thalamus begins prenatally and continues postnatally (22, 23). Glu, the primary excitatory amino acid, plays a major role in the maturing neuronal circuitry (24, 25). Glu increases significantly and drives the observed increase in Glx, since changes in Gln are not significant. The rapid increase in these neuronal metabolites likely reflects the underlying changes in growth, myelination and neurotransmission which are enabled by the increase in oxygenation and oxidative metabolism during early postnatal life (26, 27). This increase is consistent with the expected increases in energy metabolism and neurotransmission during postnatal life. tCr includes contributions from creatine and phosphocreatine, a phosphate buffer critical to rapid ATP replenishment (2) in high energy demanding central nervous system catalyzed by creatine kinase activity (28), and shows a significant increase from HERCULES. PRESS-derived mI levels decrease significantly, in agreement with previous reports (3, 4, 29). mI, a carbocyclic sugar localized to the glial compartment, has been previously shown to promote synaptic connectivity in early postnatal brain development (30).

We did not observe significant linear effects of age for any lower-concentration metabolites. GABA, the predominant inhibitory neurotransmitter, has highest concentration in DGN, including the thalamus. GSH, the most abundant antioxidant, has highest concentrations in cortex and periventricular regions. Other lower-concentration metabolites measured include Asc (Vitamin C), a neuroprotective antioxidant (31); NAAG, a neuromodulator more concentrated in WM than GM (32, 33); and PE, a key lipid precursor (34). HBCD is the first large-scale study to investigate low-concentration metabolites across infancy and childhood, and it remains to be determined what the age-related trajectories will look like across a larger timescale. Prior studies have shown that GABA+ is larger in adolescents compared to infants (35), so we expect an age-related increase across longer time spans. By contrast, much of what is known about GSH in infants has been abstracted from preclinical models. GSH in postnatal rat brain development is low initially and peaks during a period of synaptogenesis (36). Therefore, we expect that GSH will increase with age. Asc significantly decreases in rat pups during the postnatal period (37). Postnatal 7–28 days in the rat study (37) can correspond to a substantially different developmental time window in humans(38). PE concentrations in early infancy (∼6 mmol/kg) are three-fold higher than the typical values observed in adults, as observed from a ^31^P-MRS study (39).

Short-TE and HERCULES-Sum spectra are separate acquisitions with many shared features suggesting that the pattern of results should be similar across both analyses. The major singlets are prominent in both, and some signals (e.g. Glu and Gln) may be better resolved in the sparser HERCULES-Sum. Age-related correlations appeared to be stronger in the HERCULES data for tNAA, tCr, Glu and Glx than in the short-TE data, possibly due to ∼2x greater SNR. However, the stronger correlations with age observed from HERCULES measurements may also be driven by higher metabolite T_2_-weighting, shifting slopes to be more positive for all metabolites. HERCULES data are acquired at a longer TE, compared to short-TE PRESS, although the short-TE water reference is used for quantification in both cases. Osprey workflow uses adult metabolite and water T_1_ and T_2_ literature values. These might bias metabolite measurements, since these relaxation times are different from the pediatric brain as observed from 1.5T studies (29). Furthermore, pediatric relaxation values may also change across this period, influencing the observed trajectories. Therefore, future analysis with age-appropriate relaxation values for 3T is necessary. Age-appropriate tissue water density correction is also crucial in pediatric MRS quantification. HBCD QALAS (8, 11) data could provide this essential information for correction in future analyses.

It is an unresolved question whether (or under what experimental conditions) Glu and Gln are reliably separable at 3T (40). We report combined Glx as well as Glu and Gln, noting that our median tCr linewidth (∼4 Hz) is substantially less than in adults, enhancing resolution of overlapping multiplets. Signals that are not reliably resolved in the adult spectrum may be resolvable in these data. Interestingly, the findings from both datasets are the same – Glx increases with age, and that increase is driven by Glu. This supports the idea that Glu and Gln can be resolved in short-TE spectra with excellent SNR and linewidth.

The linewidth exclusion criterion applied here is stringent, perhaps beyond required levels, but there are resolution/modeling benefits to a linewidth criterion below typical coupling constants (7 Hz), and conservatism is merited for this analysis of tabulated results without individual review of spectra. This cut-off does not represent a precedent/recommendation for future analyses, particularly ones that integrate various data quality metrics. Some poorly-modeled PRESS datasets, excluded with high relRes, had excellent linewidth and SNR – it should be possible to retrieve these data in future analyses. The basis sets with which Osprey models the spectra have a linewidth of 1 Hz, not much less than the Lorentz component of the narrowest lines; narrower lines are also harder to locate and much less tolerant of frequency shifts. Many relRes exclusions of HERCULES spectra arise from subtraction artefacts and poor frequency alignment of narrow spectra. These aspects of Osprey pre-processing and modeling may be improved in future releases; this dataset will inform those developments.

One limitation of this study pertains to participant “ages”. To maintain participant anonymity, “chronological age” in the tabular data release is jittered by up to 7 days, so we did not use that noisier measure. Instead, we utilized adjusted age, which is rounded down to the nearest week, thus quantized, rather than being a continuous variable, resulting in an ‘x-axis’ age error (of up to 1 week). Future analyses of Release 2.0 with a wider age range will mitigate this error, as well as the use of the imaging file-based metadata, which includes a more precise measure of participant age. Although these data span 0–10 weeks of adjusted age, this range is not uniformly sampled – Visit-2 protocol aims for 0–1 month imaging, with flexibility for convenience, resulting in fewer datapoints from 5–10 weeks. This distribution limits power to reveal cross-sectional correlations in this dataset, but power will improve when including additional visits.

Overall, this dataset shows large variability in concentration levels at each age arising from: individual neurometabolic differences independent of age; individual differences in neurometabolic development; variability associated with rounding of the age variable; fundamental measurement variance; instrumental variance across sites including field drift, and many other sources. A previous study (4), exploring non-linear trajectories of neurometabolite levels in multiple regions, showed more variability in the DGN than in WM regions or parietal/occipital GM.

For edited MRS, editing efficiency is closely linked to field stability. Real-time frequency drift correction using the interleaved water references is implemented within the Philips sequence(9), but not yet on Siemens/GE. Harmonization techniques like ComBat (41) or inclusion of site covariates in future analyses could help address site-induced variability. At this stage, we did not exclude datapoints based on environmental exposures. Data collection is ongoing and HBCD plans annual releases in February each year. Release 2.0 will include neuroimaging data from Visits 2, 3, and 4, enabling longitudinal analyses of neurometabolite trajectories and non-linear trajectories of the metabolites of interest.

## List of Abbreviations

Asp: Aspartate
Cho: Choline
Cr: Creatine
DGN: Deep gray nuclei
GABA: Gamma-aminobutyric acid Glu Glutamate
Gln: Glutamine
Glx: Glutamine + Glutamate
GSH: Glutathione
NAA: N-Acetylaspartate
NAAG: N-Acetrylaspartylglutamate
MRS: Magnetic Resonance Spectroscopy
MRI: Magnetic Resonance Imaging
mI: Myo-inositol
sI: Scyllo-inositol
PE: Phosphorylethanolamine
PCr: Phosphocreatine
PRESS: Point-resolved Spectroscopy
HERCULES: Hadamard Editing Resolves Chemicals Using Linear-combination Estimation of Spectra
ISTHMUS: Integrated short-TE and Hadamard-edited multi-sequence
HBCD: HEALthy Brain and Child Development
TE: Echo time
tNAA: total N-Acetylaspartate
tCr: total creatine
tCho: total choline
TR: Repetition time
PCA: post-conceptional age
fMRI: Functional MRI
dMRI: Diffusion MRI
relRes: Ratio of the normalized sum of squares of the residual to the variance of the spectral noise

