## Supplementary Material for "Age-related differences in neurometabolite concentrations during early infancy: A cross-sectional analysis of the HBCD Release 1.0"

**Supplementary Table 1: MRSinMRS[11] Data Acquisition Parameters**

| **Table S1: Acquisition** | |
|---|---|
| a. Pulse sequence | ISTHMUS |
| b. Volume of Interest (VOI) | Bilateral thalamus |
| c. Nominal VOI size | 30 (RL) × 23 × 23 $mm^3$ |
| d. Repetition time (TR) | 2000 ms |
| e. Echo time (TE) | Dual TE:<br>Short-TE: 35 ms<br>Long TE: 80 ms |
| f. Total number of transients | 32 PRESS, 224 HERCULES,<br>4 Short-TE water scans, 4 long-TE water scans |
| g. Additional sequence parameters | |
| h. Water suppression method | - |
| i. Shimming method, reference peak | - |
| j. Trigger or motion correction | No trigger or active motion correction |

**Supplementary Table 2: MRSinMRS[11] Data Analysis Methods and Output**

| **Table S2: Data Analysis Methods and Output** | |
|---|---|
| a. Analysis Software | Osprey 2.8.1 |
| b. Processing steps deviating from Osprey | Segmentation in BIBSNET;<br>Localizer registration in BIDS App 2.4.3 |
| c. Output measure | Tissue-corrected water-scaled metabolite levels |
| d. Quantification references and assumptions, fitting model assumptions | Basis set list: Asc, Asp, Cr, CrCH2, GABA, GPC, GSH, Gln, Glu, mI, Lac, NAA, NAAG, PCh, PCr, PE, sI, Tau, MM09, MM12, MM14, MM17, MM20, Lip09, Lip13, Lip20<br>Fitting method: Osprey baseline knot spacing 0.4 ppm |
